# Derivation of stationary distributions of biochemical reaction networks via structure transformation

**DOI:** 10.1101/2021.03.23.436681

**Authors:** Hyukpyo Hong, Jinsu Kim, M Ali Al-Radhawi, Eduardo D. Sontag, Jae Kyoung Kim

## Abstract

Long-term behaviors of biochemical reaction networks (BRNs) are described by steady states in deterministic models and stationary distributions in stochastic models. Unlike deterministic steady states, stationary distributions capturing inherent fluctuations of reactions are extremely difficult to derive analytically due to the curse of dimensionality. Here, we develop a method to derive analytic stationary distributions from deterministic steady states by transforming BRNs to have a special dynamic property, called complex balancing. Specifically, we merge nodes and edges of BRNs to match in- and out-flows of each node. This allows us to derive the stationary distributions of a large class of BRNs, including autophosphorylation networks of EGFR, PAK1, and Aurora B kinase and a genetic toggle switch. This reveals the unique properties of their stochastic dynamics such as robustness, sensitivity and multi-modality. Importantly, we provide a user-friendly computational package, CASTANET, that automatically derives symbolic expressions of the stationary distributions of BRNs to understand their long-term stochasticity.

## Introduction

A standard approach to mathematical modeling of biochemical reaction networks (BRNs) is to use ordinary differential equations (ODEs), whose variables represent concentrations of molecules (*1*). However, this deterministic description, while convenient for computation, by its nature cannot capture the inherent randomness of BRNs. In particular, the long-term behavior of ODE systems is characterized by steady states or other attractors, rather than by the stationary distributions statistically observed in real biological systems. As cell biology moves away from bulk averages to single-cell measurements, a focus has shifted to the study of such stationary distributions (*2, 3*). They can be described by various stochastic approaches (*1, 4*). In particular, stationary distributions can be described as steady-state solutions of the chemical master equation (CME), which has been widely used to describe the time evolution of the probabilities for the numbers of chemical species in BRNs such as gene regulatory networks and signaling pathways (*5*).

Since the CME is a differential equation with infinitely many variables, its steady-state solution (i.e., the stationary distribution) can be found analytically only for simple cases, such as linear reaction networks (*6*) or birth-death processes (*7*). Unlike the CME, its deterministic counterpart is a finite dimensional ODE, whose steady-state solutions are relatively easier to calculate. An interesting question, therefore, is whether there is a systematic way of using these deterministic steady states for characterizing the stationary distribution of the stochastic counterpart. There is a positive answer to this question for special networks, called complex balanced networks.

A result from queuing theory (*8*), reinterpreted in the context of BRNs (*9*) through the connection between Petri nets and BRNs (*10*), shows that for complex balanced networks whose kinetics are described by mass action reactions, stationary distributions can be characterized in terms of jointly distributed Poisson random variables with parameters corresponding to deterministic steady states. An independent proof of this result, together with deep applications to CMEs, was developed by Anderson, Craciun, and Kurtz (*11*). Complex balancing is difficult to check and depends on rate constant values. However, beautiful work by Horn, Jackson, and Feinberg (*12–14*) has shown that all networks that have the special structural properties of weak reversibility and zero deficiency are complex balanced, independently of rate constants. Weak reversibility of a network means that the network is a union of closed reaction cycles, and the deficiency of a network is the number of dependent closed reaction cycles, which can be easily checked. Satisfying these two structural properties is a simple condition to derive the stationary distribution of network under mass action reactions with the method in (*11*).

As various BRNs such as networks of several reversible reactions (e.g., *A* + *B* ↔ *C* ↔ 0) or cyclic reactions (e.g., *A* → *B* → *C* → *A*) are weakly reversible and deficiency zero, their stationary distributions can be analytically derived (*15–20*). These have been used to characterize the stochasticity of various systems, including a genetic oscillator (*21*) and a competitive inhibition enzyme kinetics model (*22*). Unfortunately, the majority of BRNs do not have the special network structure. For instance, only ~0.36% of the Erdős-Rényi random networks of two species with up to bimolecular reactions have a deficiency of zero when the edge probability is 0parameter values are set as follows:.5, and the fraction decreases to zero as the number of species increases (*23*). Moreover, from a biological standpoint, even simple networks are unlikely to be weakly reversible if they include a bimolecular reaction whose reverse reaction is unimolecular (e.g., autophosphorylation and dephosphorylation).

Here, we develop a framework to derive stationary distributions for a class of networks which do not have the special structure (i.e, weakly reversibility and zero deficiency) by modifying their structures via *network translation* (*24, 25*). Specifically, by simply merging reactions with a common stoichiometric vector and shifting reactions in the networks, we are able to change their structure to be weakly reversible and deficiency zero while preserving their stochastic dynamics. This allows us to derive the stationary distributions of a large class of BRNs including autophosphorylation networks of EGFR, PAK1, and Aurora B kinase. This derivation reveals key reactions determining the autophosphorylation status, which can seldom be done with a purely numerical approach. Furthermore, we describe how the stochastic dynamics of more complex BRNs can be tracked when our method is applicable for only their subnetworks. Importantly, we provide a user-friendly computational package CASTANET (Computational package for deriving Analytical STAtionary distributions of biochemical reaction networks with NEtwork Translation) that automatically derives the stationary distribution of submitted BRNs via our method. This will provide an effective tool to analyze the stochasticity of BRNs.

## Results

### Obtaining the desired network structure via network translation

As mentioned in the introduction, the stationary distributions of the stochastic mass action models for BRNs can be derived with any choice of rate constants using the method in (*11*) if and only if the networks have two structural properties: *weak reversibility* and *zero deficiency.* However, even very simple networks such as the one shown in Fig. 1a-left fail to satisfy the two properties. Weak reversibility means that if there exists a path from a *complex* (i.e., a node in the reaction graph) to another complex, then there is a reverse path from the second one back to the first one. Because there is no path from *A* + *A* to *A* while there is a path from *A* to *A* + *A*, the network in Fig. 1a-left is not weakly reversible. The deficiency of a network is a non-negative integer index calculated by subtracting both the number of linkage classes (i.e., connected components in the reaction graph) and the dimension of the subspace spanned by the stoichiometric vectors from the number of complexes. The deficiency of the network in Fig. 1a-left is one. Therefore, the method in (*11*) cannot be used to derive its stationary distribution.

**Fig. 1:**
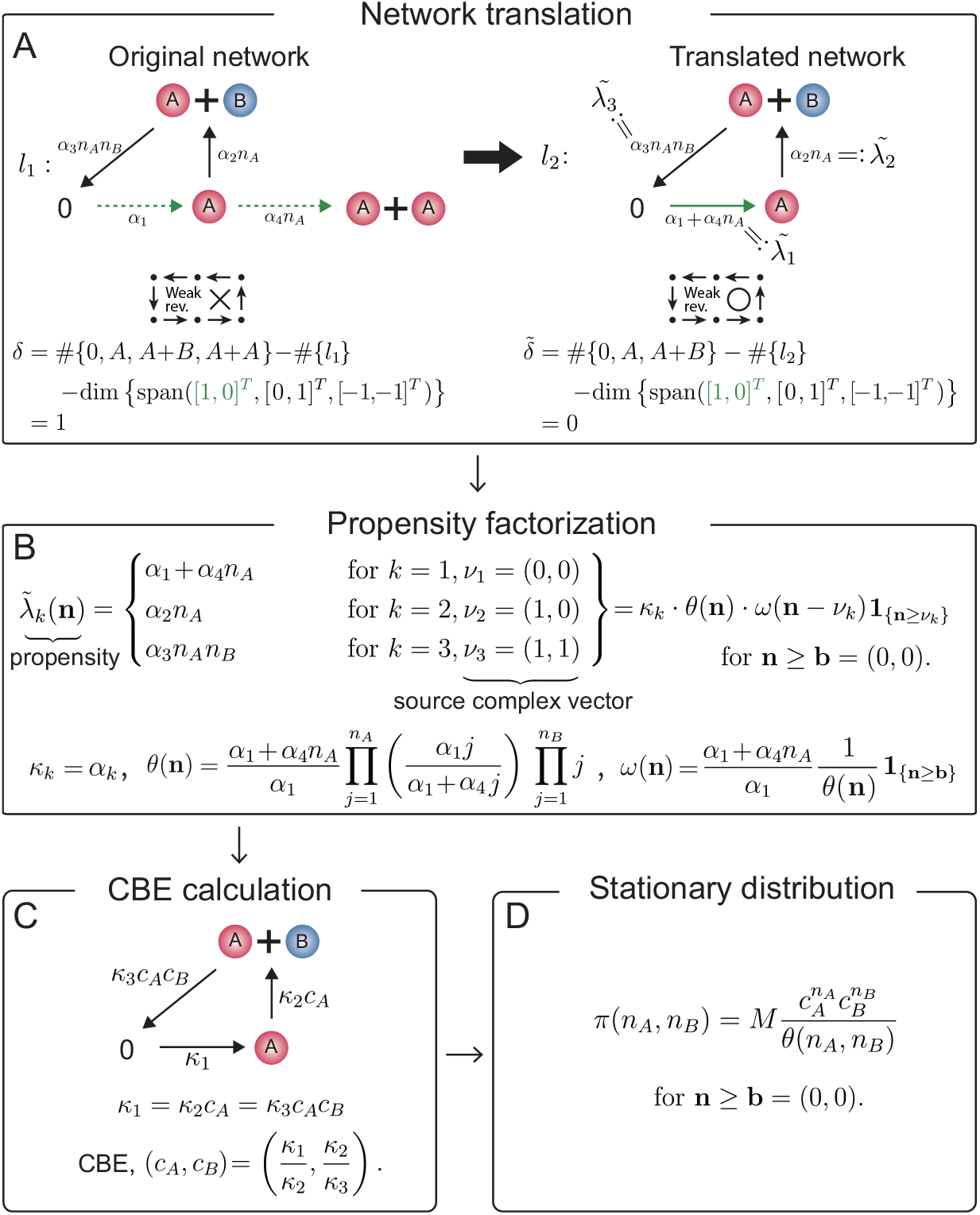
Derivation of a stationary distribution with network translation. **a** The non-weakly reversible and deficiency (*δ*) one network is translated to the weakly reversible deficiency zero network by merging two reactions, which have the same stoichiometric vectors (green dotted lines). 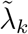 denotes the propensities of the translated network. **b** Factorize 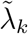 with constants *κ*_*k*_ and functions *θ*(**n**) and *ω*(**n**) as 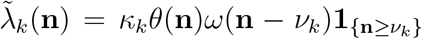 on Γ = {**n** | **n ≥ b**} at which 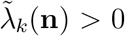 if **n** ≥ *ν*_*k*_ + **b**, and 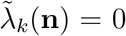 otherwise. *ν*_*k*_ is the source complex vector of the *k*th reaction. **c** Compute a CBE of the deterministic mass action model for the translated network with rate constants {*κ*_*k*_}. **d** Using the *θ*(**n**) and the CBE, the stationary distribution can be derived analytically. Here, *M* is a normalizing constant.

Two different reactions, 0 → *A* and *A* → *A* + *A*, have the same stoichiometric vector (1, 0) because both reactions produce one molecule of *A* (Fig. 1a-left). Thus, these two reactions can be merged by unifying the source complexes 0 and *A* into 0 and summing the propensities of both reactions (Fig. 1a). This procedure is known as *network translation* (*24, 25*), which was proposed to investigate deterministic systems. This procedure is also applicable to stochastic systems as it preserves the stochastic dynamics (see Supplementary information for details). For instance, the propensities of the production of *A* are *α*_1_ + *α*_4_*n*_*A*_ in both the original (Fig. 1a-left) and the translated network (Fig. 1a-right). Although the network translation is simple, it can effectively change the structure of the network to be a weakly reversible deficiency zero network.

### Propensity factorization is required

Even though the translated network is weakly reversible and of zero deficiency, the new model no longer follows mass action kinetics since the propensity of the reaction 0 → *A* is not constant (Fig. 1a-right). In this case, previously, it was known that the method in (*11, 26*) is still applicable if the non-mass action propensity functions can be factorized as in a certain form. However, the propensity functions of this translated network do not have the certain form. Thus, we generalize the previous factorization form so that stationary distributions can be derived for a larger class of BRNs. Specifically, we show that all the propensities of the translated network 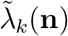 need to be factorized as

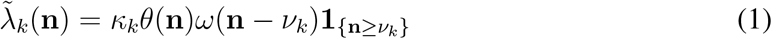

for some constants *κ*_*k*_ > 0 and functions *θ*(**n**) > 0 and *ω*(**n**) ≥ 0 on a set Γ = {**n** | **n** ≥ **b**} where the *ν*_*k*_ is the source complex vector of the *k*th reaction, the inequality is coordinate-wise, and the **b** needs to be chosen so that 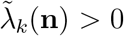 if and only if there are sufficient reactants (i.e., **n** ≥ *ν*_*k*_ + **b**) in Γ. For the translated network (Fig. 1a-right), 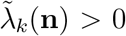 if and only if **n** ≥ *ν*_*k*_ like mass action kinetics, and thus **b** = (0, 0) and 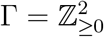.

Propensity functions satisfying the factorization condition (Eq. (8)) include the following generalized mass action kinetics (Eq. (7)). For instance, if a source complex is 0, *A, A* + *A,* or *A* + *B*, propensity functions following the generalized mass action kinetics are proportional to 1, *f*_*A*_(*n*_*A*_), *f*_*A*_(*n*_*A*_)*f*_*A*_(*n*_*A*_ − 1), or *f*_*A*_(*n*_*A*_)*f*_*B*_(*n*_*B*_), respectively. Note that if the *f*_*i*_’s are identity functions then the propensities follow standard mass action kinetics (Eq. (6)) (*11, 27*). The propensity functions following the generalized mass action kinetics can be easily factorized with 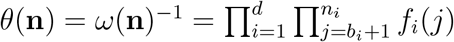, where *d* is the number of the constitutive chemical species (see Eq. (9) for details). However, the translated network (Fig. 1a) does not follow the generalized mass action kinetics (Eq. (7)) because the propensity function of the reaction 0 → *A*, *α*_1_ +*α*_4_*n*_*A*_, is not proportional to 1 (i.e., it is not constant). Thus, we need to solve recurrence relations as described in Supplementary information to identify the propensity factorization (Fig. 1b):

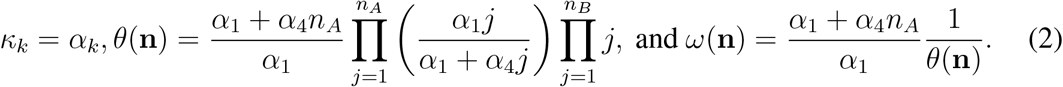

### Derivation of stationary distribution

After identifying *κ*_*k*_, *θ*(**n**), and *ω*(**n**) via the propensity factorization, we need to find a complex balanced equilibrium (CBE) of the deterministic mass action model with rate constants {*κ*_*k*_} for the translated network (Fig. 1c). The CBE is a steady state at which for each complex *ν*, the in-flow rate to *ν* is equal to the out-flow rate from *ν* (*12*). For instance, based on the deterministic model in Fig. 1c, the complex balance conditions for the complex 0, *A*, and *A* + *B* are *κ*_3_*c*_*A*_*c*_*B*_ = *κ*_1_, *κ*_1_ = *κ*_2_*c*_*A*_, and *κ*_2_*c*_*A*_ = *κ*_3_*c*_*A*_*c*_*B*_, respectively. By solving these equations, we can obtain the CBE, (*c*_*A*_, *c*_*B*_) = (*κ*_1_/*κ*_2_, *κ*_2_/*κ*_3_). Note that the existence of a CBE is guaranteed because we translate a network to be weakly reversible and deficiency zero (*13*).

Finally, using the function *θ*(**n**) (Fig. 1b) and the CBE (*c*_*A*_, *c*_*B*_) (Fig. 1c), we can derive the stationary distribution of the stochastic model for the translated network, which is the same as that of the original network, as follows:

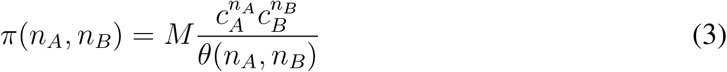

for *n*_*A*_ ≥ 0, *n*_*B*_ ≥ 0 where *M* is the normalizing constant so that the sum of the stationary distribution is one (see Methods for details). In this example, the distribution *π*(**n**) is obtained on 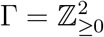. This state space is closed as proved by Theorem 2 in Supplementary information, and it is irreducible (i.e., every state is reachable from every other state; see Supplementary Remark 4 for details). On the other hand, if an irreducible state space is a proper subset of Γ, possibly due to a conservation law, then the normalizing constant *M* is chosen so that the sum of *π*(**n**) over the subset is one.

### Computational package, CASTANET

Applying our theoretical framework (Fig. 1) has two practical difficulties. Translating a given network to a weakly reversible deficiency zero network (Fig. 1a) is not straightforward as prohibitively many candidates of translated networks often exist. Furthermore, it is challenging to check whether the factorization condition holds (Fig. 1b) as it requires to solve associated recurrence relations. Thus, we have developed a user-friendly, open-source, and publicly available computational package, “CASTANET (https://github.com/Mathbiomed/CASTANET),” that automatically performs network translation and propensity factorization and derives stationary distributions (Fig. 2a). With this package, we were able to easily identify hundreds of BRNs and derive analytic forms of their stationary distributions. We have provided some of them in Fig. 2b and Supplementary Figs. 3 and 4. To use this package, users only need to enter the source complexes, product complexes, and propensity functions of reactions.

**Fig. 2:**
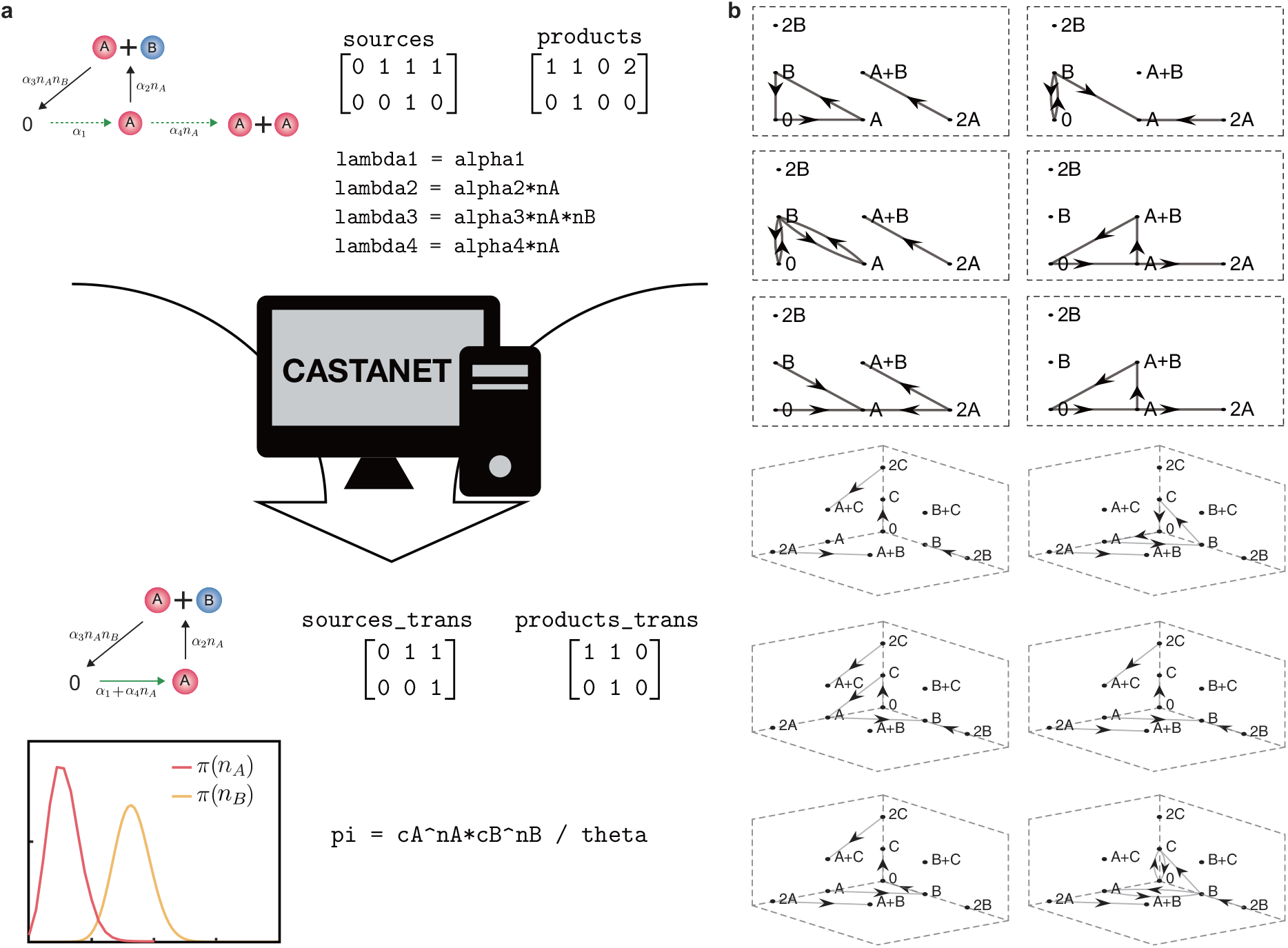
CASTANET (Computational package for deriving Analytical STAtionary distributions of biochemical reaction networks with NEtwork Translation). **a** A schematic diagram for the computational package. If users simply enter the source complexes, product complexes, and propensity functions of reactions (`lambda_k`), then the package identifies a weakly reversible deficiency zero translated BRN (`sources_trans and products_trans`) and then derives its stationary distribution (`pi`). See Supplementary information for a step-by-step manual. **b** BRNs with two species (top) and three species (bottom) whose stationary distributions were calculated by our computational package. The tail and head of each arrow represent the source and product complexes of reactions, respectively. They are assumed to follow the stochastic mass-action kinetics, and the rate constants can take any positive values. See Supplementary Figs. 3 and 4 for more examples. Here, each network is embedded in euclidean spaces where we present A+A and B+B as 2A and 2B, respectively.

### Stationary distributions of autophosphorylation networks

Our theoretical framework and especially CASTANET extend the class of BRNs whose stationary distributions can be derived analytically using CBEs (Figs. 1 and 2). This class includes various autophosphorylations networks (Fig. 3) that are not weakly reversible due to autophosphorylation reactions, which occur in intermolecular (trans), intramolecular (cis), or mixed manners (*28, 29*).

**Fig. 3:**
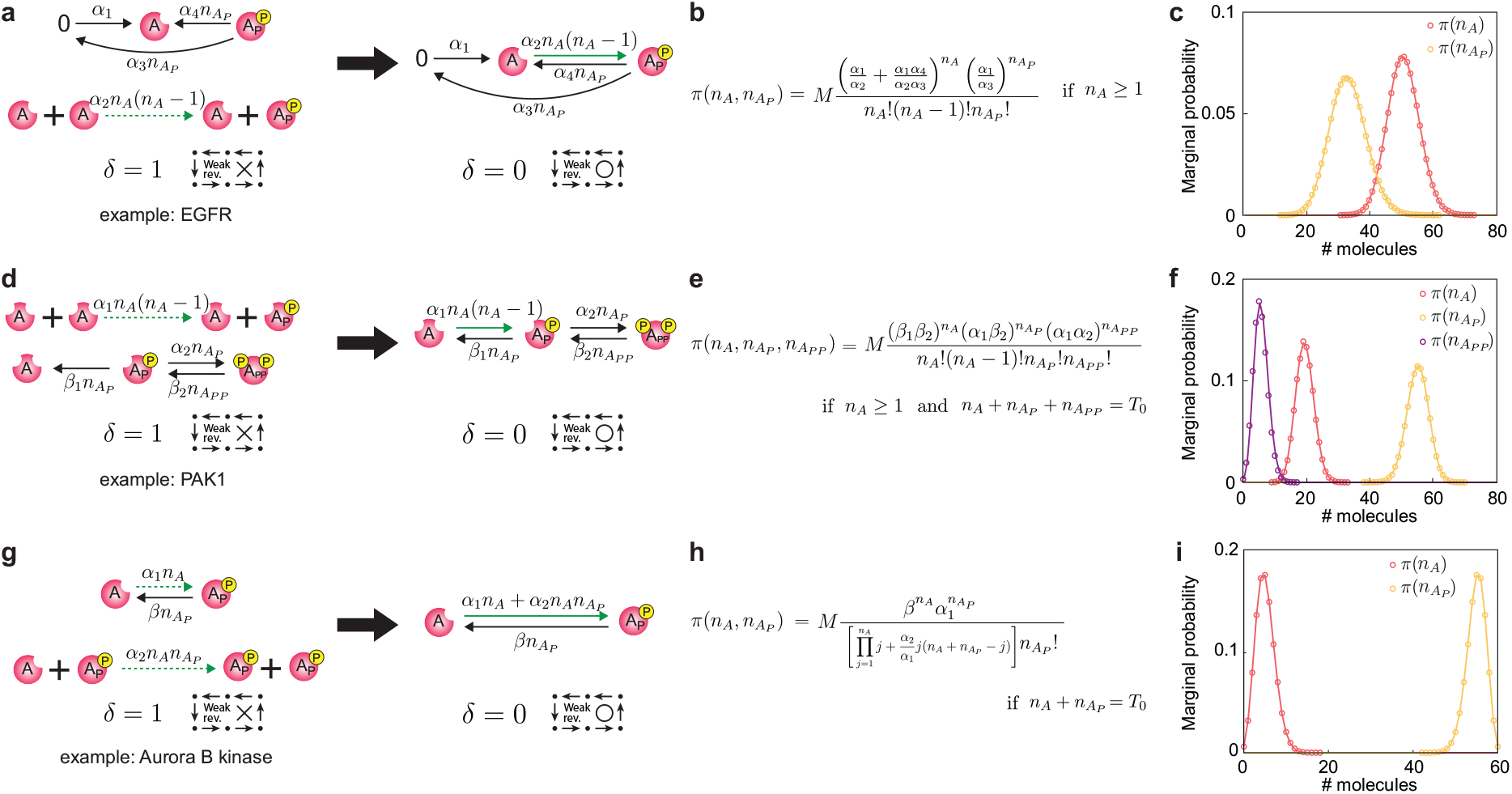
Stationary distributions of diverse autophosphorylation networks. **a, d, g** While the autophosphorylation networks have a deficiency of one and are not weakly reversible, they can be translated to weakly reversible deficiency zero networks. **b, e, h** T hus, stationary joint distributions can be derived using the method illustrated in Fig. 1. *T*_0_ in E and H represent the total numbers of proteins, which are conserved. **c, f, i** The marginal probabilities of the numbers of species derived from the formula (solid line) and stochastic simulations (dot) are consistent. Here, parameter values are set as follows: **a** *α*_1_ = 10, *α*_2_ = 0.03, *α*_3_ = 0.3, *α*_4_ = 2, **d** *α*_1_ = 0.3, *α*_2_ = 0.1, *β*_1_ = 2, *β*_2_ = 1, *T*_0_ = 80, **g** *α*_1_ = 0.001, *α*_2_ = 1, *β* = 5, *T*_0_ = 60. For each example, 10^5^ simulations were performed using the Gillespie algorithm (*30*).

Asymmetric trans-autophosphorylation occurs if two monomers form a homodimer and one of them acts as an ‘enzyme’ and phosphorylates the other. This type of autophosphorylation occurs in the epidermal growth factor receptor (EGFR), which triggers signal transduction for cell proliferation (*31*). The key regulatory reactions for EGFR include its synthesis, trans-autophosphorylation, dephosphorylation, and degradation (Fig. 3a-left). The asymmetric trans-autophosphorylation is a reaction that transforms the complex *A*+*A* to *A*+*A*_*P*_. The dephosphorylation reaction is not the reverse of the previous reaction; instead, it occurs from the complex *A*_*P*_ to the complex *A*. Thus, the network is not weakly reversible. However, CASTANET automatically identifies a weakly reversible deficiency zero translated network and its propensity factorization (Fig. 3a-right) and then derives the analytic form of stationary distribution (Fig. 3b) that matches what is calculated with stochastic simulations (Fig. 3c).

In addition, having the formula (Fig. 3b) allows us to easily understand the long-term behavior of the system, something that is not possible with a purely computational approach. For instance, 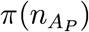 in Fig. 3c is the Poisson distribution with rate *α*_1_/*α*_3_. This indicates that the synthesis (*α*_1_) and degradation rates (*α*_3_) of *A* are the determinants of the long-term status of 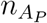, which is surprisingly robust to the changes of phosphorylation (*α*_2_) and dephosphorylation rates (*α*_4_). Furthermore, *π*(*n*_*A*_) is solely determined by 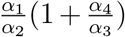, and its moments can be calculated with the modified Bessel functions (see Supplementary information for details). This allows us to identify that the stationary distribution of *n*_*A*_ is sub-Poissonian, and its coefficient of variation attains the maximum at 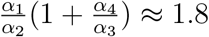 (Supplementary information Fig. 1).

Trans- and cis-autophosphorylation can occur sequentially. For example, p21-activated kinase 1 (PAK1), which regulates cell motility and morphology, phosphorylates a threonine residue in the kinase domain in a trans manner asymmetrically (*A* + *A* → *A* + *A*_*P*_) (*28, 32*), and then phosphorylates a serine residue in the regulatory domain of itself in a cis manner (*A*_*P*_ → *A*_*PP*_) (*33, 34*) (Fig. 3d-left). While the original network of PAK1 is not weakly reversible (Fig. 3d-left), CASTANET identifies a weakly reversible deficiency zero translated network (Fig. 3d-right).and derives the analytic form of stationary distribution (Fig. 3e) that matches the simulation result (Fig. 3f).

Both trans- and cis-autophosphorylation can occur simultaneously as in Aurora B kinase, which controls mitotic progression (*35*). In an Aurora B kinase network, cis-autophosphorylation (*A* → *A*_*P*_) promotes rapid trans-autophosphorylation (*A* + *A*_*P*_ → *A*_*P*_ + *A*_*P*_), which forms a positive feedback in the system (*35*). For this network, CASTANET successfully applies our method to derive the analytic form of stationary distribution (Fig. 3g-i).

While mass action kinetics are commonly used to describe autophosphorylations (*35, 36*) as in our examples, the Michaelis-Menten function and Hill function are also often used (*37, 38*). Moreover, they are also used to describe the effects of phosphatases on dephosphorylation and proteasomes on degradation (*39, 40*), which EGFR, PAK1, and Aurora B kinase undergo (*41–46*). Even when the mass action propensities in the networks (Fig. 3a, d, and g-left) are replaced with the Michaelis-Menten or Hill functions, their stationary distributions can still be derived with the same approach (Supplementary Fig. 5).

When the presented networks are extended by adding reactions, our methods might not be applicable. For instance, if an additional trans-autophosphorylation (*A* + *A*_*P*_ → *A*_*P*_ + *A*_*P*_) is added to the example in Fig. 3a, although it can be translated to a weakly reversible deficiency zero network, their propensities cannot be factorized as in Eq. (1). Thus, the stationary distribution of the extended network cannot be derived by our method. However, it can be approximated by the stationary distribution of the original network if the rate constant of the added reaction is small enough (see Supplementary Fig. 6 for details). Such approximation works for the extended networks of the other networks (Fig. 3d, g) as well. This indicates that if the stationary distributions of core subnetworks, which consist of dominant reactions, can be derived by our method, then it could be used to approximate the stationary distributions of their more complex parent networks.

### Translation of fast subnetworks reveals both the fast and slow dynamics of a multi-timescale system

As the number of nodes (i.e. species) of networks increases, the networks are less likely to be a weakly reversible deficiency zero network even after network translation, and thus our method is less likely to be applicable. However, such large networks commonly consist of reactions occurring at different time scales (*47*). In this case, if we can derive the conditional stationary distributions of only fast subnetworks with our method, both the fast and slow dynamics of the full network can be accurately captured.

For gene regulatory networks, if the promoter kinetics (i.e., binding and unbinding of transcription factors to promoters) are fast, the fast subnetwork is a simple reversible binding network (i.e., weakly reversible and of zero deficiency), and thus its stationary distribution can be easily calculated (*21*). On the other hand, when the promoter kinetics are slow, the fast subnetwork includes a complex protein reaction network whose stationary distribution is challenging to derive. This can occur for a variety of reasons, e.g., the presence of nucleosomes in eukaryotic cells usually slows down the binding and unbinding of transcription factors (*20*).

A genetic toggle switch with the slow promoter kinetics, which consists of a pair of genes *G*^*A*^ and *G*^*B*^, is an example of such multi-timescale system (Fig. 4a). The genes *G*^*A*^ and *G*^*B*^ express proteins *A* and *B*, respectively. Subsequently, these proteins undergo asymmetric trans-autophosphorylation, and they mutually repress gene expression by binding to the promoter region of each other’s gene. The binding and unbinding of the phosphorylated proteins occur at a slower time scale. Therefore, the entire network can be divided into the fast subnetwork consisting of gene expression, phosphorylation, dephosphorylation, and degradation and the slow subnetwork consisting of binding and unbinding (Fig. 4b). While the fast subnetworks of *A* and *B* are neither weakly reversible nor deficiency zero, they can be translated to weakly reversible deficiency zero networks (Fig. 4c-top). The propensity functions of these translated networks can be factorized as in Eq. (8), so their stationary distributions conditioned on the slow variables can be derived with CASTANET (Fig. 4c). Depending on the slowly changing gene states, the distributions of the proteins *A*, *A*_*P*_, *B*, and *B*_*P*_ can dramatically change.

**Fig. 4:**
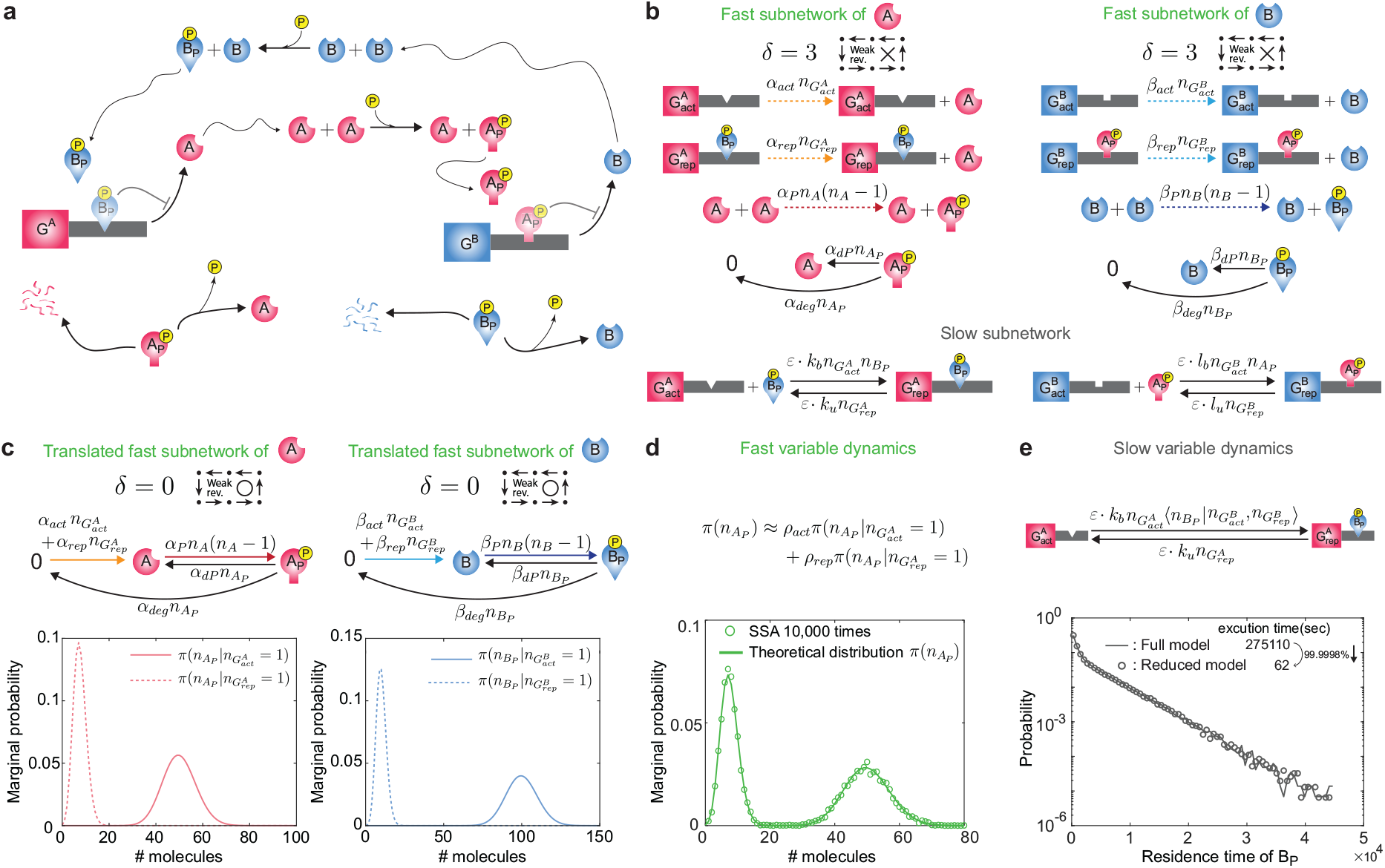
Fast and slow dynamics of a multi-timescale system are identified via network translation. **a** Schematic diagram of a toggle switch system. **b** Protein kinetics are fast but promoter kinetics (activation and repression of the genes) are slow. The fast subnetworks of *A* and *B* are not weakly reversible and have deficiencies of three. **c** Translated fast subnetworks, obtained by merging reactions having the same stoichiometric vectors (colored arrows), are weakly reversible and deficiency zero. Stationary distributions of the number of the phosphorylated proteins conditioned on the gene states (activated/repressed) are derived by using the method illustrated in Fig. 1. **d** The unconditional stationary distribution of fast variables, which is bimodal, can be accurately approximated by the weighted average of the conditional stationary distributions. The weights *ρ*_*act*_ and *ρ*_*rep*_ are the probabilities that the gene *G*^*A*^ is active and repressed, respectively. **e** By replacing the fast variable 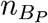 with its conditional stationary moment 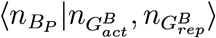 determined by the current state of the slow variables 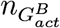 and 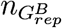, the reduced model can be derived, which can capture the slow dynamics of the genes. For instance, the reduced model (dots) accurately captures the residence time distributions of the repressor *B*_*P*_ to its target gene *G*^*A*^ of the full model (solid line). The execution times for performing 10^4^ simulations with the full and the reduced models are 275110 and 62 seconds, respectively. Parameter values are set as: *ϵ* = 10^−5^, *α*_*act*_ = 10, *α*_*rep*_ = 1.5, *α*_*P*_ = 0.2, *α*_*dP*_ = 1, *α*_*deg*_ = 0.2, *β*_*act*_ = 10, *β*_*rep*_ = 1, *β*_*P*_ = 0.3, *β*_*dP*_ = 2, *β*_*deg*_ = 0.1, *k*_*b*_ = 1, *k*_*u*_ = 30, *l*_*b*_ = 1.3, and *l*_*u*_ = 20.

When the gene states slowly change, the fast variables rapidly equilibrate to the conditional stationary distributions determined by the current gene states (Fig. 4c). Thus, the weighted average of the conditional stationary distributions with the probabilities of the corresponding gene state accurately approximates the full (i.e., unconditional) stationary distribution under timescale separation (i.e., *ε* ≪ 1) (*20*). For instance, the full stationary distribution of *A*_*P*_ can be approximated as

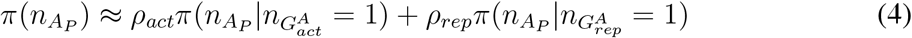

where *ρ*_*act*_ and *ρ*_*rep*_ are the probabilities that the gene *G*^*A*^ is active and repressed, respectively. The *ρ*_*act*_ becomes larger as the dissociation constant between *G*^*A*^ and its repressor *B*_*P*_ is larger, and the number of repressor *B*_*P*_ is smaller. The *ρ*_*act*_ can be calculated by identifying the eigen-vector of the matrix consisting of the dissociation constant, and the conditional stationary moments of the repressors obtained from Fig. 4c (*20*), and *ρ*_*rep*_ = 1 − *ρ*_*act*_ (see Supplementary information for details). Thus, using Eq. (4), we can accurately capture the bimodal stationary distribution of the protein *A*_*P*_ (Fig. 4d), leading to phenotypic heterogeneity in isogenic populations. Similarly, the full bimodal stationary distributions of the other fast variables *A, B,* and *B*_*P*_ can also be accurately captured (Supplementary Fig. 7). Note that these bimodalities cannot be captured by the corresponding deterministic model, which predicted monostability (the result is not shown). Such mismatches between the stochastic and deterministic model have been frequently observed in the presence of timescale separation (*1, 20, 48*).

The conditional stationary distributions of the fast variables, obtained by using our approach (Fig. 4c), allow us to capture the slow dynamics of the full system as well. On the slow time scale, the slow variables are unlikely to be changed, but the fast variables rapidly equilibrate to their conditional stationary distributions for the given slow variables. Thus, by replacing the fast variables in the propensity functions of the slow reactions with their quasi-steady states (QSSs): conditional stationary moments, we can obtain the reduced model (*21, 49*). For the toggle switch system, the QSSs of the fast variables *A*_*P*_ and *B*_*P*_ can be computed from their conditional stationary distributions (Fig. 4c). Then by replacing the fast variables *A*_*P*_ and *B*_*P*_ with their QSSs, we can obtain the reduced model with only the slow variables, the active and repressed genes (Fig. 4e). This reduced model accurately captures the slow dynamics of the full model: the binding and unbinding of the repressors to the genes. Both the full and the reduced models yield nearly identical distributions of the residence time of the repressor *B*_*P*_, which quantifies how long the repressor maintains its binding to the gene *G*^*A*^ (Fig. 4e). Because the reduced model does not simulate the fast reactions, which incur a large computational cost in the full model simulation, computation time decreases by 99.9998%.

## Discussion

In this study, we have developed a framework and its computational package that analytically derive stationary distributions of a large class of BRNs. Specifically, we showed that the stationary distribution of a BRN can be derived if two conditions are satisfied: the network can be transformed to a weakly reversible deficiency zero network via network translation (Fig. 1a) and the propensity functions of the translated network satisfies the generalized factorization property of mass action kinetics, identified in this study (Fig. 1b). We found that these conditions are satisfied in numerous BRNs including various autophosphoryaltion networks by using CASTANET (Fig. 2, and Supplementary Figs. 3, 4). Furthermore, even when only a subnetwork of more complex BRNs satisfies the conditions, the stochastic dynamics can often be captured. That is, the stationary distribution of the subnetwork consisting of dominant reactions derived with our method can accurately approximate the stationary distribution of its parent network (Supplementary Fig. 6). Furthermore, the derivation of the stationary distribution of a fast subnetwork is enough to capture both the slow and fast stochastic dynamics of its multi-timescale parent network (Fig. 4). With these analytically derived stationary distributions of BRNs, their long-term stochastic behaviors such as their dependence on rate constants can be effectively investigated, and the likelihood function of parameters for Bayesian inference can also be derived (*2*).

Our work focused on the derivation of steady-state solutions of the CME using the underlying network structure following previous studies (*11, 26*). However, the CME is not usually used to capture cell division, which should be taken into account to describe single cell behavior in general. Thus, it would be interesting in future work to extend our method to the population balance equation (*50, 51*), which describes stochastic cell population dynamics (e.g. cell division) as well as intracellular dynamics. This extension could be accomplished by averaging stationary distributions from cell populations after a stationary distribution of each cell is derived by our method.

We have translated a network to have the desired structural properties (i.e., weak reversibility and zero deficiency) by merging reactions with a common stoichiometric vector (Figs. 1a and 3g) and shifting a reaction preserving its stoichiometric vector (Fig. 3a, d). While the idea underlying this procedure is simple, it greatly extends the class of networks whose structure can be changed to the desired one. For instance, when the edge probability is 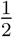, the fraction of deficiency zero networks among Erdős-Rényi random networks with two species and at most bimolecular reactions increases more than six times after network translation. The identification of such translation, which is not simple, can be done automatically by the provided computational package, CASTANET. In particular, to efficiently search translated networks, in CASTANET, we use the necessary conditions for network translation toward weakly reversible and deficiency zero networks, derived in this study (Theorems 3 and 4 in Supplementary information).

Furthermore, CASTNET performs the propensity factorization of the translated networks, which is required to derive the stationary distributions of networks with non-mass action kinetics. In this study, by extending the previous factorization condition (*11, 26*) to ours (Eq. (8)), we have been able to derive stationary distributions of various BRNs (Figs. 1, 2, and 3, Supplementary Figs. 3 and 4). Although the factorization condition with non-mass action kinetics have been rarely investigated (*11, 26*) due to its complexity and lack of motivation, our work motivates studies on it as translated networks typically follow non-mass action kinetics. To cover more weakly reversible deficiency zero translated networks, we aim to further generalize our factorization conditions, and accordingly, we will update our computational package CASTANET.

By changing the network structure while preserving the stochastic dynamics via network translation, we have been able to use the theory, applicable to weakly reversible deficiency zero networks (*11*), to understand the stochastic dynamics of a larger class of networks with non-zero deficiencies. Similarly, by translating networks to have a deficiency of one, it would be possible to show that the networks have the properties of a network with a deficiency of one, such as absolute concentration robustness: the steady state value of a species is invariant to the overall input of the system (*52–54*). Furthermore, network translation of stochastic BRNs can also be used to identify stochastic properties of networks based on their structures, such as positive recurrence (*55*) and extinction (*53, 56, 57*).

## Methods

### Biochemical reaction network

BRN is a graphical representation of a given biochemical system (*12, 14, 58, 59*). It consists of the triple 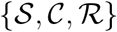 where 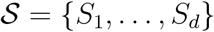 is the set of interacting *species*, 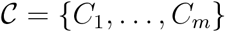 is the set of *complexes*, and 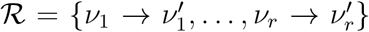 is the set of *reactions*. A complex is a non-negative linear combination of species (i.e., *C*_*i*_ = *a*_*i*1_*S*_1_ + ⋯ + *a*_*id*_*S*_*d*_), which is also represented as a *d*-dimensional non-negative integer-valued vector (*a*_*i*1_, … , *a*_*d*_). A reaction is an ordered pair of complexes. This allows the BRN to be represented as a directed graph 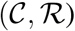, where complexes are the nodes and reactions are directed edges. Hence, a reaction 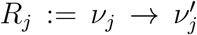, where *ν*_*j*_ and 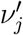 are the *source* and *product* complexes of the *j*th reaction, respectively. The vector 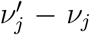 is called a *stoichiometric vector* of the *j*th reaction, which describes the relative change in the number of molecules of reactants and products between the sides of each reaction. A *linkage class* is a connected component of the network when all reactions are regarded as undirected edges. Weak reversibility means that if there is a sequence of reactions from a complex *C*_*i*_ to another complex *C*_*j*_ then there must be a sequence of reactions from *C*_*j*_ to *C*_*i*_. The deficiency *δ* is the integer index defined as 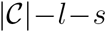, where 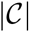 is the number of complexes, *l* is the number of linkage classes, and *s* is the dimension of the subspace spanned by all stoichiometric vectors (Fig. 1a).

### Complex balanced equilibrium

CBE of the deterministic mass action model for a BRN with rate constants {*κ*_*k*_} is the steady state 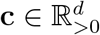, which satisfies the following equality for each complex 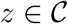 (Fig. 1c):

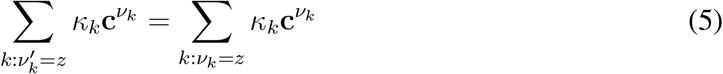

where the 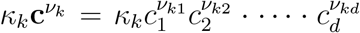 is the rate function of the *k*th reaction following the deterministic mass action kinetics, and *ν*_*ki*_ is the *i*th entry of *ν*_*k*_ (*12*). The LHS is the sum of rate functions over reactions whose product complex is *z* (i.e., 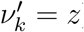), and the RHS is the sum of rate functions over reactions whose source complex is *z* (i.e., *ν*_*k*_ = *z*). In other words, at CBE, the in-and out-flows create a balance for each complex. The deterministic mass action model for a BRN possesses a CBE regardless of rate constants if and only if the BRN is weakly reversible and deficiency zero (*13*). Furthermore, even when a BRN has non-zero deficiency and is weakly reversible, the deterministic mass action model for the BRN possesses a CBE with specific choice of rate constants (*13*).

### Stochastic model of biochemical reaction networks

We model a BRN as a continuous-time Markov chain (CTMC) for an isothermal well-stirred system with constant volume. The state of the CTMC at time *t*, 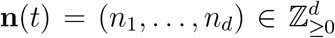, represents the copy number of each species. Each reaction is associated with a *propensity function*:

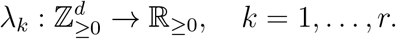

Specifically, *λ*_*k*_(**n**) is the probability that the *k*th reaction occurs in a short interval of length *dt* if the state at the beginning of the interval was **n**. Using the propensity functions, we can derive the CME, which describes the time evolution of the probability of the model:

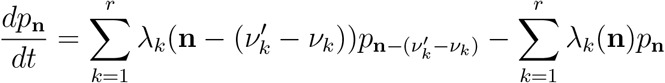

for 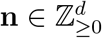, where *p*_**n**_(*t*) denotes the probability that the state of the system equals 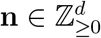 at time *t*. A *stationary distribution π*(**n**) of a given CTMC is the steady-state solution of the CME that satisfies the following infinite equation:

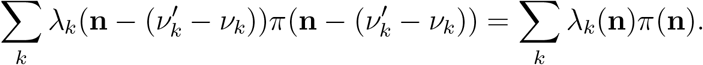

It means that if the CTMC is initialized with its stationary distribution, the vector of probabilities *p*(*t*) will stay constant for all time *t* > 0.

The stochastic mass action propensity functions are assumed to be proportional to the number of ways in which species can combine to form the source complex. Hence, the *k*th propensity function with the rate constant *α*_*k*_ can be written as:

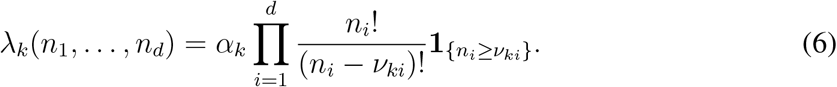

Additionally, the propensity functions can have the more generalized form as follows:

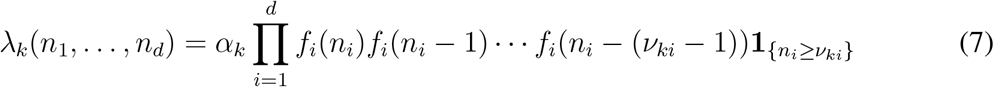

where functions *f*_*i*_ : ℤ_≥0_ → ℝ_≥0_. For instance, the translated network in Fig. 3a-right follows this form as *f*_*A*_(*n*_*A*_) = *n*_*A*_(*n*_*A*_ − 1) and 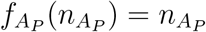. This is called ‘generalized’ stochastic mass action kinetics since if *f*_*i*_’s are identity functions then it is equivalent to the stochastic mass action kinetics (Eq. 6).

### Network translation

Network translation is a procedure to transform a BRN 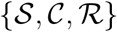 to another one 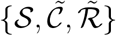 that satisfies the condition: the sum of propensities of a set of reactions sharing the same stoichiometric vector remains identical (Fig. 1a). That is, for each vector *γ* ∈ ℤ^*d*^, the propensity functions of the original and the translated networks, *λ*_*k*_(**n**) and 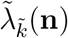, satisfy the following:

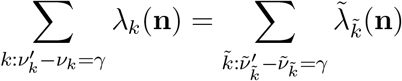

for all 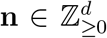, where 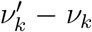 and 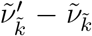 are the stoichiometric vectors of the *k*th and 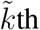 reactions of the first and second models, respectively. For example, merging several reactions sharing a common stoichiometric vector into one reaction is network translation. Similarly, shifting reactions preserving their stoichiometric vectors is also an instance of network translation (e.g., *A* + *B* → 2*B* to *A* → *B*). Network translation can change the structural properties of BRNs, such as weak reversibility and the deficiency, but it preserves the associated CME, i.e, stochastic dynamics (see Supplementary information for details).

### Propensity factorization

To derive the stationary distribution with our approach (Fig. 1), all the propensities 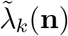 should be factorized as

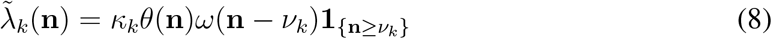

for some constants *κ*_*k*_ > 0 and functions *θ*(**n**) > 0 and *ω*(**n**) ≥ 0 on a set 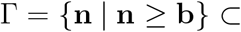 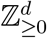 at which 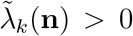 if **n** ≥ *ν*_*k*_ + **b** and 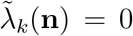 otherwise. For the stochastic mass action kinetics (Eq. (6)), **b** = **0** as 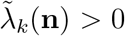 if and only if **n** ≥ *ν*_*k*_. For the translated network in Fig. 3a, **b** = (1, 0) because each propensity is positive if and only if **n** ≥ *ν*_*k*_ + (1, 0) in 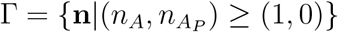.

While the propensity factorization can be calculated by solving recurrence relations (see Supplementary information for details), it can be obtained without solving recurrence relations if all the propensities of a given network follow the generalized mass action kinetics (Eq. (7)).

In this case, the factorization can be easily obtained as

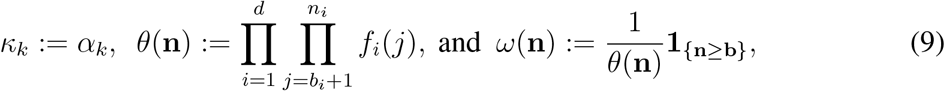

where 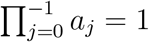 for any {*a*_*j*_} (see Supplementary information for details).

### Derivation of stationary distributions

If a network is weakly reversible and deficiency zero so that it has a CBE 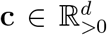 (*13*) and propensity function *λ*_*k*_(**n**) can be factorized as in Eq. (8) on Γ, a stationary measure of the network can be derived as

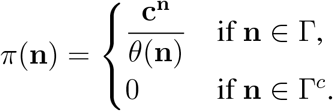

Supplementary information provides the proof and detailed illustration. By scaling this stationary measure with the normalizing constant, which is the reciprocal of the summation of *π*(**n**) over the irreducible state space, the stationary distribution on the irreducible state space can be obtained. For instance, the normalizing constant for the stationary distribution (Fig. 3e) is calculated by summing *π*(**n**) over the irreducible state space 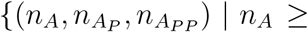 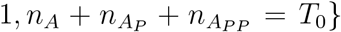. While computing the normalizing constants is sometimes challenging, a symbolic computation approach using Wilf-Zeilberger theory can be used for the stochastic mass action model (*60*).

### Computational package, CASTANET

We have developed a user-friendly, open-source, and publicly available computational package, CASTANET, that performs the network translation and propensity factorization automatically (Fig. 2a). The package checks two conditions: whether a given BRN can be made weakly reversible and of zero deficiency after network translation, and whether the propensities of the translated network can be factorized as in Eq. (8). If these two conditions are satisfied, CASTANET then calculates the analytic formula for a stationary distribution.

To efficiently search weakly reversible deficiency zero translated networks, we derived their necessary conditions (Theorems 3 and 4 in Supplementary information) and incorporated them in the package. Furthermore, CASTANET constructs a candidate for the factorization function *θ*(**n**) in symbolic expression, which allows us to check propensity factorization condition without checking infinite combinations (see Supplementary information).

## Data availability

All data used in the current research are available upon request to the corresponding authors.

## Code availability

The MATLAB code performing network translation, propensity factorization, and CBE calculation to derive stationary distribution (schematically shown in Figs. 1 and 2) can be found at https://github.com/Mathbiomed/CASTANET. The detailed description and step-by-step manual are provided in Supplementary information.

## Acknowledgments

This work is supported by the Samsung Science and Technology Foundation (SSTF-BA1902-01)(JKK) and the National Research Foundation of Korea (Global Ph D. Fellowship Program 2019H1A2A1075303)(HH), as well as US NSF grants 1716623 and 1849588 (EDS). JK thanks German Enciso for travel support via NSF grant DMS1616233. HH and JK thank the 2019 Chemical Reaction Networks summer school and workshop at Politecnico di Torino for providing valuable feedbacks on this work.

## Author contributions

HH, EDS and JKK designed research. HH, JK, and JKK developed the theory and algorithm. HH performed computation. All authors discussed and analyzed results. HH, JK, and JKK drafted the manuscript and designed the figures, and all authors revised the paper.

## Supplementary information

Supplementary Text Figs. S1–S7

Supplementary References

